# Ten complete mitochondrial genomes of Gymnocharacini (Stethaprioninae, Characiformes): evolutionary relationships and a repetitive element in the Control Region (D-loop)

**DOI:** 10.1101/2020.10.26.355867

**Authors:** Rubens Pasa, Fabiano Bezerra Menegídio, Igor Henrique Rodrigues-Oliveira, Iuri Batista da Silva, Matheus Lewi Cruz Bonaccorsi de Campos, Dinaíza Abadia Rocha-Reis, John Seymour (Pat) Heslop-Harrison, Trude Schwarzacher, Karine Frehner Kavalco

## Abstract

We are presenting the complete mitogenomes of eight fish species/cytotypes from Neotropical region belonging to the *Astyanax* and *Psalidodon* genus: *A. aeneus, A. altiparanae, P. fasciatus* (from two locations - Upper Paraná and São Francisco river basins), *A. lacustris, P. rivularis* (two cytotypes) and *P. rioparanaibano*. We perform the whole-genome sequencing for six of these species in a Novaseq 6000 - by Illumina, meanwhile two genomes were assembled from raw data available in databases. Plus, we reassembled and annotated the mitochondrial genomes for *A. mexicanus* and *P. paranae*, both already described and with raw data available online. All the genomes presented the same organization, with 13 protein-coding genes, 22 tRNA genes and two rRNA genes. Aiming to contribute to the understanding of the several cryptic species complexes and phylogeny of the genus, we perform Bayesian analysis using the 13 protein-coding genes from these species, plus *Deuterodon giton* and using a *Brycon* species as outgroup.

## 1. Introduction

Stethaprioninae is a subfamily of characiform fish that comprises small animals popularly known as tetras. *Astyanax* was the first genus in the number of species within the subfamily, being the most diverse in the Neotropical region [1]. Historically, the genus has a wide geographical distribution, through the south USA to the north of Argentina [2], but some *Astyanax* species share several common features that difficult their identification and recognition [3] leading to efforts to identify diagnostic characteristics or molecular signatures for the group. Recently, the genus *Astyanax* underwent an initial taxonomic review and several of its species have been reclassified in new or disused (old) genera [4]. Some of the species formerly identified as belonging to *Astyanax* were reassigned as *Psalidodon* or *Deuterodon*, for example. Terán et al [4] moved some species of *Astyanax* to others six genera, including the species complexes *A. fasciatus* [5] and *A. scabripinnis* [6] (relocated to *Psalidodon*) and the *Astyanax* species from Coastal river basins of Brazil, like as *A. giton* (relocated to the genus *Deuterodon)*, while the species of the complex *A. bimaculatus* [7] and the North American species remain in the *Astyanax* genus [4]. Especially for the tetras from South America, there is a good background of information from chromosomal and phylogenetic data that indicates trends in the evolution of this group [8].

Despite the great diversity and the taxonomical issues, at least one species of the genus (*A. mexicanus*) have been used as a model to understand the development of eyes and the evolution of complex traits [9] [10] [11]. This species was named *A. fasciatus* in the past, with epigean and hypogean (troglomorphic) morphotypes. However, currently, *P. fasciatus* is a designation restrict to tetras from Brazilian rivers and probably corresponds to a group of cryptic species with the largest diversity in diploid numbers in *Psalidodon* genus, with 2n=45, 46, 47, 48 and 49 chromosomes, plus the presence of B chromosomes and heterochromatin polymorphisms in some populations [12] [13] [14], within low molecular diversification [15]. Morphologically, the populations of the São Francisco river basin are different from the specimens from upper Paraná River and Paraíba do Sul [16] and the original morphological description is so broad that it certainly covers other species outside the complex [17]. In this case, *P. fasciatus* should be only the specimens from the São Francisco river basin (original basin of the type species), and the others may be cryptic species of the complex or even other species [17]. On the other side, despite a clear genetic structure between populations from Upper Paraná and São Francisco river basins, morphometric traits seem to be homoplasy [18].

The *A. bimaculatus* complex is currently represented by the homonymous species, along with others such as *A. altiparanae* (considered as a junior synonym of *A. lacustris* by some authors), *A. lacustris, A. assuncionensis* and *A. abramis*. With a preference for calm waters, these species habits mainly in the upper Paraná, Paraguay, Iguassu and São Francisco river basins [19]. Contrary to what is observed for in *P. fasciatus* and *P. scabripinnis, A. bimaculatus* shows a constant diploidy number in different populations, 2n = 50 chromosomes, considered a symplesiomorphic character in Gymnocharacini [20] [21] [22]. The diversity in the group refers to differences in its karyotypic formula, fundamental number and general symmetry of the karyotypes [20] [22]. These cytogenetic data, associated with the molecular ones, suggest relatively recent divergence, as well as the monophyletic status of this branch [8].

The third species complex explored in this work, *P. scabripinnis*, was proposed by Moreira-Filho & Bertollo [6] based on morphological and chromosomal characteristics of specimens collected in the Paraná and São Francisco river basins. In a review of *P. scabripinnis* group, Bertaco & Lucena [23] pointed out the existence of 15 species, including *P. paranae* and *P. rivularis*. The species of this complex are known for their wide karyotypic diversity, with diploid numbers ranging from 2n=46 to 50 chromosomes [6] [24] [25]. In recent studies using molecular phylogeography and geometric morphometry, Rocha et al. [26] reinforced the validity of *P. rivularis* and *P. paranae*, sister species of the complex inhabiting the São Francisco and Paraná river basins, respectively. However, among the populations from Upper Rio Paranaíba, the existence of a new species of the complex was observed due to morphometric and mtDNA data [27]. This new species, called *Psalidodon rioparanaibano*, was collected only in a small tributary of the Paranaíba river, surrounded by populations of *P. paranae*. Moreover, within *P. paranae* and *P. rivularis* groups, karyotypic diversity is also present [28] (Rodrigues-Oliveira et al., in preparation), indicating that even though delimited by individual lineages, these groups still constitute compilations of cryptic species.

Hundreds of papers describing mitogenomes have been published in the last few years, and despite being the most sequenced genome nowadays [29], to this date, just the mitogenomes of *A. mexicanus* [30]*, P. paranae* [31], and *Deuterodon giton* [32], *P. fasciatus* and *A. altiparanae* [33] are published. In an attempt to fill this gap, we present now the complete sequences of the mitochondrial genome of ten species/cytotypes of *Astyanax/Psalidodon* genus, discussing the evolutionary relationships among these species and the presence of a repeat in the control region.

## 2. Material and Methods

We analyzed specimens belonging to the *Astyanax/Psalidodon* genus. We collected the samples in different locations through the major Brazilian rivers and their vouchers are deposited in the ichthyological collection of the Laboratory of Ecological and Evolutionary Genetics at the Federal University of Viçosa, campus Rio Paranaíba, Brazil (Table 1). After sampling, we brought the living specimens to the laboratory, euthanized them according to the ethical standards of CONCEA - National Council for Control of Animal Experimentation of Brazil and CEUA/UFV - Animal Use Ethics Committee/Federal University of Viçosa (760/2018). We performed the sampling with licenses provided by SISBIO/ICMBIO - Biodiversity Authorization and Information System (1938128) and SISGEN - National System for the Management of Genetic Heritage and Associated Traditional Knowledge (A9FE946).

**Table 1.** Ecological data from the collected specimens for this study.

| Species | Sequence | Sex | Country/basin | Lat/Lon | Altitude | Specimen voucher | Tissue type |
| --- | --- | --- | --- | --- | --- | --- | --- |
| *Astyanax altiparanae* | Mitochondrial | Male | Brazil: Paranaíba river basin | -19.351761111111<br>-46.793002777778 | 843 m | LAGEEVO - 4121 | liver and heart |
| *Astyanax lacustris* | Mitochondrial | Male | Brazil: Abaeté river basin | -20.123172222222<br>-45.970891666667 | 708 m | LAGEEVO - 3732 | liver and heart |
| *Psalidodon fasciatus* | Mitochondrial | Female | Brazil: Bambuí river basin | -19.209913888889<br>-46.109386111111 | 1013 m | LAGEEVO - 2096 | liver and heart |
| *Psalidodon rioparanaibano* | Mitochondrial | Male | Brazil: Paranaíba river basin | -19.187713888889<br>-46.236177777778 | 1074 m | LAGEEVO - 4285 | liver and heart |
| *Psalidodon rivularis* 2n = 46 | Mitochondrial | Male | Brazil: Abaeté river basin | -18.9428<br>-45.938388888889 | 889 m | LAGEEVO - 2030 | liver and heart |
| *Psalidodon rivularis* 2n = 50 | Mitochondrial | Male | Brazil: Abaeté river basin | -19.209913888888<br>-46.110219444444 | 1017 m | LAGEEVO - 2614 | liver and heart |

We extracted the total genomic DNA of six specimens *(Psalidodon rivularis* with 46 chromosomes, *Psalidodon rivularis* with 50 chromosomes, *Psalidodon fasciatus* from São Francisco river basin*, Psalidodon rioparanaibano, Astyanax lacustris*, and *Astyanax altiparanae)* from the liver and heart tissue according to the instructions of the Invitrogen’s PureLink DNA extraction and purification kit. After quality checking using fluorometer Qubit (Thermo Fisher Scientific), the Whole Genome Sequencing was performed in a Novaseq 6000 (Illumina, San Diego, CA) at Novogene company, UK.

For broader comparisons, we also assembled the mitogenome of two species with raw reads available on ENA (European Nucleotide Archive): *P. fasciatus* from Upper Parana river basin (SRR8476332) and *A. aeneus* from Mexico (SRR1927238). Aiming to validate our methodology we reassembled the mitogenomes of *P. paranae* (SRR5461470) and *A. mexicanus* (SRR2040423). In the Bayesian analysis, we also included the mitochondrial complete sequence of *Deuterodon giton* (NC_044970.1), and *Brycon orbignianus* (KY825192.1) as outgroup.

We assembled the mitogenome from raw reads on Novoplasty v3.7 [34] in a parallel cluster computer (64 Gb RAM) using the mitogenome of *Psalidodon paranae* available on GenBank (SRR5461470) as seed (supplement). We annotated the obtained sequences on MitoAnnotator [35] at MitoFish (http://mitofish.aori.u-tokyo.ac.jp). We uploaded the 2×150 raw reads to the Galaxy [36] public platform at usegalaxy.eu, where accessed the quality of raw reads (using FastQC, Babraham Bioinformatics) and filtered with Fastp tool [37].

We perform comparative genomics analysis by BLAST comparison of all *Astyanax/Psalidodon* mitochondrial genomes against a reference (*Psalidodon paranae*) generated by Blast Ring Image Generator (BRIG) [38]. To access the repetitive region we analyze the mitochondrial sequences with Tandem Repeats Finder [39], and aligned on ClustalW [40] to found the repeat motif.

We aligned Fasta sequences with ClustalW [40] and calculated the p-distance with MEGA X software [41]. We used the 13 protein-coding genes (PCGs) in Bayesian phylogenetic inference with MrBayes 3.2.7 [42] after calculation of best evolutionary models for each segment with Partition Finder 2.1.1 [43]. Bayesian analyses were performed using four independent chains with 10-million-generations and posteriorly was verifying the effective sample size (ESS) and strand convergence in Tracer 1.7 software [44]. The first 25% of the generations were discarded as burn-in. For the Maximum Likelihood analysis, we have used the concatenated 13 PCGs after the test of the best model in MEGA X software [41].

## 3. Results

Our results have shown that all mitogenomes content and gene order were identical (Figure 1), with 13 protein-coding genes (PCGs), 22 tRNA genes and two rRNA genes with the order expected according to already described Characiformes mitogenomes, including to other *Astyanax/Psalidodon/Deuterodon* species as *A. mexicanus* [30], *P. paranae* [31] and *D. giton* [32]. All PCGs, except the *Nd6* gene, are on the heavy chain. All but eight tRNAs are on the heavy chain as well.

**Figure 1:**
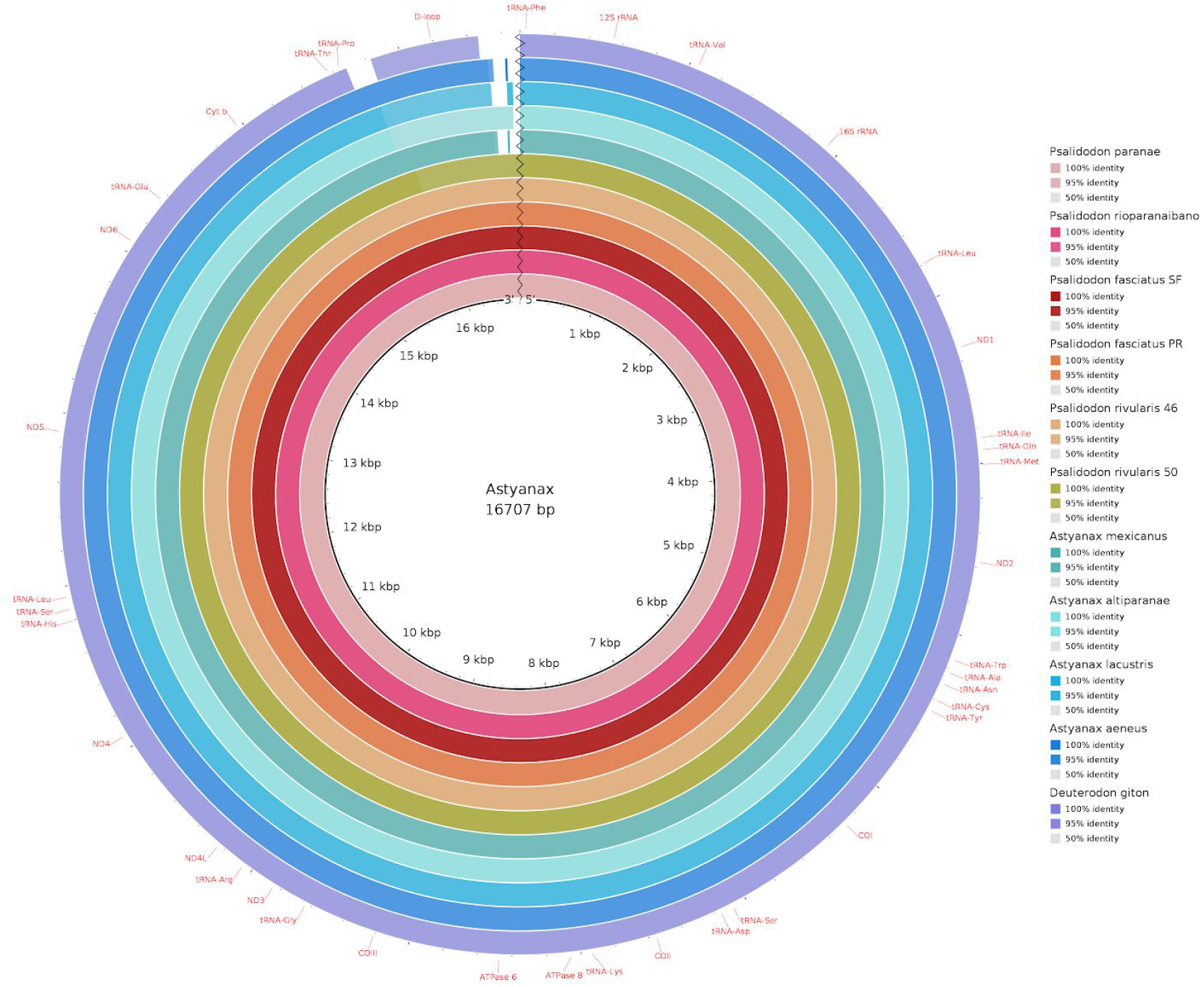
Comparative genomics analysis of all 10 Stethaprioninae fish. BLAST comparison of all mitochondrial genomes against a reference (*Astyanax paranae*) generated by Blast Ring Image Generator (BRIG). Gaps in rings correspond to regions with less than 50% identity to the reference sequence.

The statistics of the sequences are summarized in table 2. The length of mitochondrial sequence range from 16,626 bp in *Psalidodon fasciatus* from São Francisco basin, to 16,812 bp in *Psalidodon rivularis* 2n=50. The average length of D-loop was 1,061 bp. We found a repeat of 35 bp in all D-loops, except in *Deuterodon giton* (Table 3), and with alignment, we got a repeated motif (TATGTATTAGTACATATTATGCATAATTATACATA) slightly variable in some species.

**Table 2.** General statistics of the sequences analyzed. The abbreviations PR and SF after *P. fasciatus*, indicate Paranaiba river basin and São Francisco river basin respectively. The mitogenome that was not reassembled following our methodology is indicated by *.

| Species | Size<br>(bp) | %<br>GC | A | T | C | G | N | D-loop<br>(bp) | Bankit | Genbank |
| --- | --- | --- | --- | --- | --- | --- | --- | --- | --- | --- |
| *Astyanax aeneus* | 16769 | 41.1 | 5038 | 4832 | 4318 | 2581 | 0 | 1090 | BankIt2340537 | BK013055 |
| *Astyanax altiparanae* | 16730 | 41.7 | 4952 | 4795 | 4397 | 2586 | 0 | 1066 | BankIt2341301 | MT428072 |
| *Astyanax lacustris* | 16763 | 41.8 | 4959 | 4797 | 4415 | 2592 | 0 | 1099 | BankIt2340713 | MT428067 |
| *Astyanax mexicanus* | 16768 | 41 | 5046 | 4848 | 4304 | 2566 | 0 | 1091 | BankIt2341953 | BK013062 |
| *Deuterodon giton* * | 16643 | 40.7 | 4950 | 4911 | 4182 | 2600 | 0 | 976 | - | NC_044970.1 |
| *Psalidodon fasciatus PR* | 16729 | 42.5 | 4951 | 4663 | 4466 | 2647 | 0 | 1054 | BankIt2340825 | BK013061 |
| *Psalidodon fasciatus SF* | 16626 | 42.7 | 4916 | 4615 | 4460 | 2635 | 0 | 951 | BankIt2340824 | MT428071 |
| *Psalidodon paranae* | 16743 | 42.8 | 4946 | 4626 | 4526 | 2643 | 0 | 1067 | - | SRR5461470 |
| *Psalidodon rioparanaibano* | 16714 | 42.9 | 4932 | 4607 | 4526 | 2648 | 0 | 1039 | BankIt2340806 | MT428068 |
| *Psalidodon rivularis 2n=46* | 16779 | 42.9 | 4952 | 4622 | 4541 | 2663 | 0 | 1102 | BankIt2340815 | MT428069 |
| *Psalidodon rivularis 2n=50* | 16812 | 42.8 | 4972 | 4642 | 4536 | 2661 | 0 | 1136 | BankIt2340820 | MT428070 |
| Average | 16734.2 | 41 | 4964.9 | 4723.5 | 4424.6 | 2620.2 | 0 | 1061 |  |  |

**Table 3.**
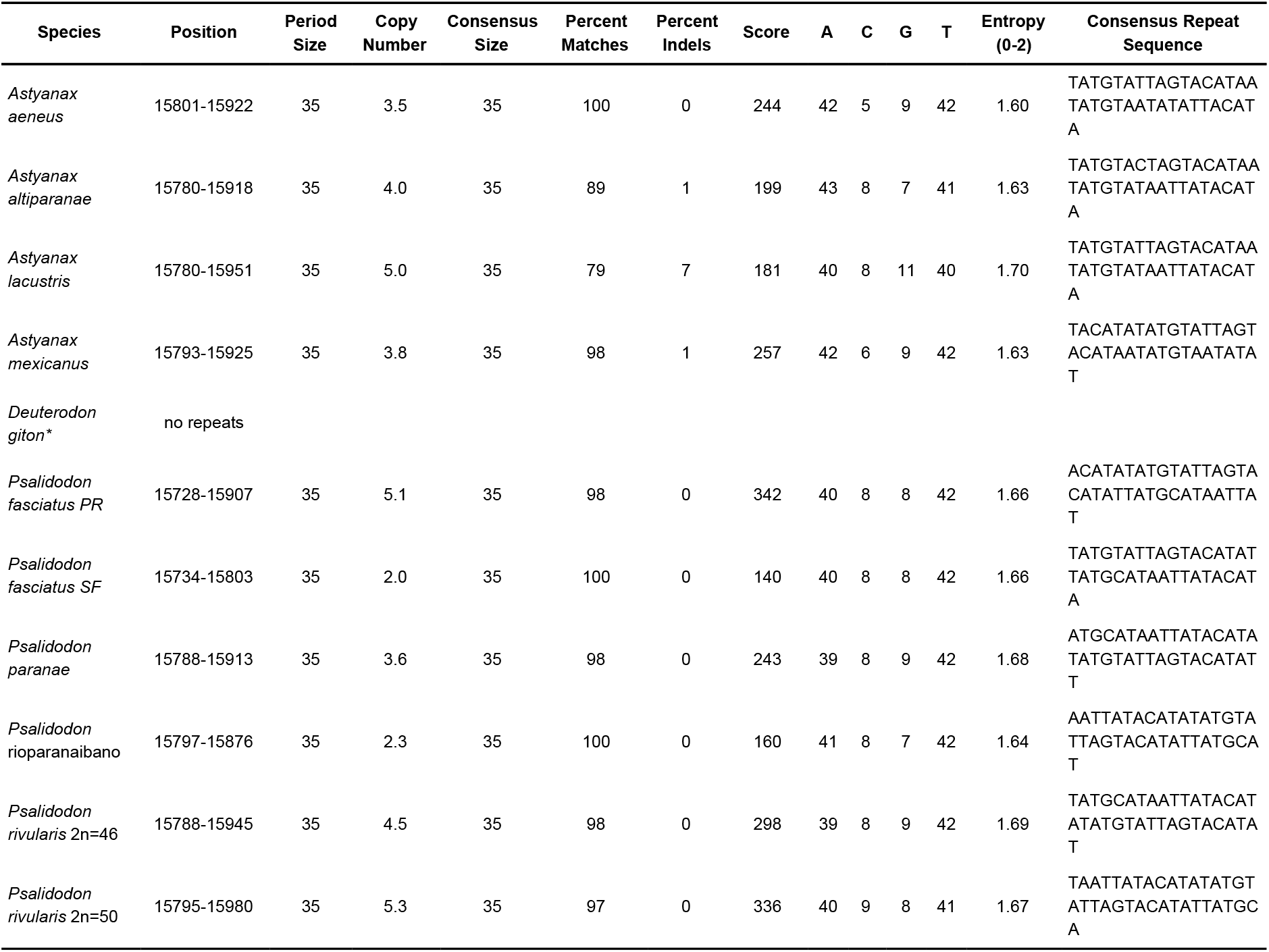
Summary of the properties from the repeated sequence found in the D-loop region. The abbreviations PR and SF after *P. fasciatus*, indicate Paranaiba river basin and São Francisco river basin respectively. The mitogenome that was not reassembled following our methodology is indicated by *.

The genetic distance among species (Table 4) is reflected in both the Maximum Likelihood and the Bayesian tree (Figure 2) that shows strong construction with high bootstrap value and posterior probabilities, respectively.

**Table 4.** Genetic distances. The abbreviations PR and SF after P. fasciatus, indicate Paranaiba river basin and São Francisco river basin respectively.

|  | *Astyan aeneus* | *Astyanax altiparanae* | *Astyanax lacustris* | *Astyanax mexicanus* | *Deuterodon giton* | *Psalidodon fasciatus PR* | *Psalidodon fasciatus SF* | *Psalidodon paranae* | *Psalidodon rioparanaibano* | *Psalidodon rivularis* 2n = 50 | *Psalidodon rivularis* 2n =46 |
| --- | --- | --- | --- | --- | --- | --- | --- | --- | --- | --- | --- |
| *Astyanax altiparanae* | 0.15159 |  |  |  |  |  |  |  |  |  |  |
| *Astyanax lacustris* | 0.15168 | 0.01802 |  |  |  |  |  |  |  |  |  |
| *Astyanax mexicanus* | 0.02153 | 0.15208 | 0.15278 |  |  |  |  |  |  |  |  |
| *Deuterodon giton* | 0.20154 | 0.20706 | 0.20697 | 0.20196 |  |  |  |  |  |  |  |
| *Psalidodon fasciatus PR* | 0.15402 | 0.16102 | 0.16128 | 0.15425 | 0.19751 |  |  |  |  |  |  |
| *Psalidodon fasciatus SF* | 0.15413 | 0.16078 | 0.16104 | 0.1541 | 0.19716 | 0.00516 |  |  |  |  |  |
| *Psalidodon paranae* | 0.14967 | 0.15824 | 0.15911 | 0.15016 | 0.1983 | 0.03719 | 0.03761 |  |  |  |  |
| *Psalidodon rioparanaibano* | 0.15107 | 0.15833 | 0.1585 | 0.15182 | 0.19725 | 0.03938 | 0.0398 | 0.02344 |  |  |  |
| *Psalidodon rivularis* 2n = 50 | 0.15089 | 0.1592 | 0.15938 | 0.15252 | 0.19795 | 0.03763 | 0.03796 | 0.02012 | 0.02161 |  |  |
| *Psalidodon rivularis* 2n =46 | 0.15072 | 0.15911 | 0.15955 | 0.15217 | 0.19883 | 0.03816 | 0.0384 | 0.02064 | 0.02318 | 0.01758 |  |
| *Brycon orbignyanus* | 0.27482 | 0.27684 | 0.27605 | 0.27579 | 0.27149 | 0.26684 | 0.26725 | 0.26566 | 0.26654 | 0.26663 | 0.26681 |

**Figure 2:**
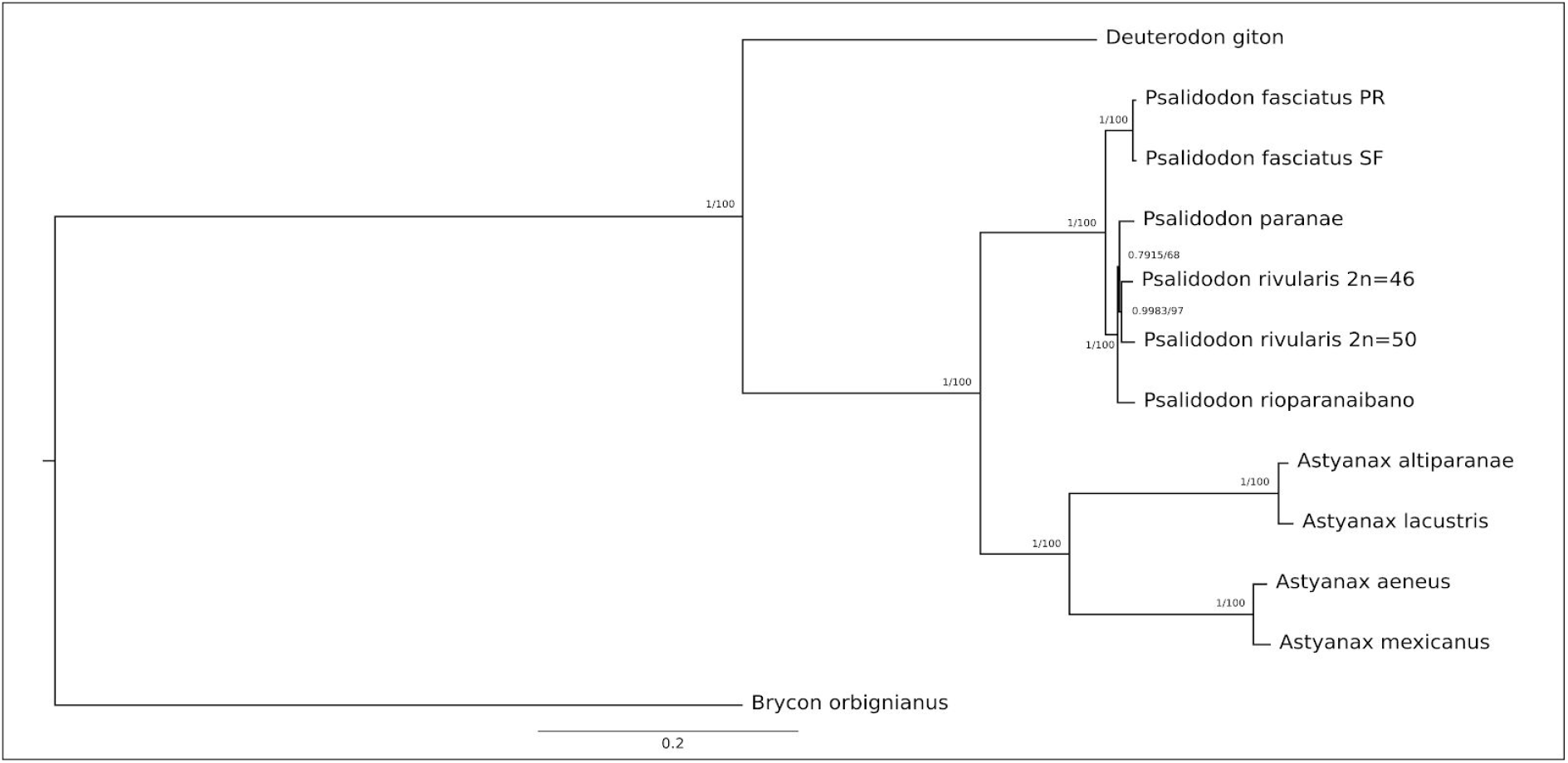
Phylogenetic tree based on 13 protein-coding genes (PCGs) showing the relationships among the Stethaprioninae fish using *Brycon orbignianus* as outgroup. The topology was the same in Maximum Likelihood (100 bootstrap replicates) after the test of the best model (General Time Reversible +G) and Bayesian Inference after calculation of best evolutionary models for each segment, using four independent chains with 10-million-generations (The first 25% of the generations were discarded as burn-in). The posterior probabilities/bootstrap are on the branches.

## 4. Discussion

The topology of the tree is congruent with those inferred for Rossini *et al*. [45] and Pazza *et al*. [8] except for the North American clade, which appear as a sister group of *A. altiparanae* and *A. lacustris* here. Besides, our study reinforces the taxonomic review by Terán et al. [4].

In this study, we observed that the populations of *P. fasciatus* belonging to the Paraná and São Francisco river basins had the lowest genetic distances (0.5%, Figure 2). The genetic distance found is similar to that obtained by Pazza et al. [18] for the mitochondrial Cytochrome Oxidase I gene (COI; 0.6%), and substantially lower than that of the mitochondrial ATPase 6/8 gene (1.9%). Thus, it is possible that among the analyzed species complexes, *P. fasciatus* is the one with the most recent adaptive radiation history. Combined with the genetic structure found between different basins [18], the high karyotype diversity present in the group [12] [13] [14] [15] reinforces a tendency to maintain different Evolutionarily Significant Units (ESUs) in each population [46].

In a review based on meristic characters, Lucena and Soares [47] consider *A. bimaculatus* and *A. altiparanae* to be a junior synonym of *A. lacustris*. Despite the monophyly of these two species (Figure 2), previously observed within the *A. bimaculatus* complex [8], the genetic distance between them (1.8%, Table 4) suggests that both units correspond to independent species isolated in different river basins. Additionally, studies of geometric morphometry indicate that the morphological similarity in this group has been shaped by parallel evolution [48], which would explain the difficulty in establishing autapomorphies for each species.

Instead, *A. altiparanae* and *A. lacustris*, which occur widely in different hydrographic basins, the two analyzed cytotypes of *P. rivularis* coexist in a small geographical area (Abaeté sub-basin river). In this scenario, the genetic distance found (1.76%, Table 4) added to the karyotypic differences (2n = 46 and 2n = 50), are strong indications that both cytotypes correspond to cryptic species with distribution limited to the Upper São Francisco river basin. It is possible that speciation and the evolution in *P. rivularis* was, or is being driven mainly by chromosomal diversification, with the subsequent genetic divergence, as observed in *P. fasciatus* [15].

Our study also reaffirmed the validity of *P. rioparanaibano* as an evolutionary significant unity (ESU) distinguishable from *P. paranae* [27], although both are found in the Paranaíba river basin. Besides the great genetic distance observed between *P. paranae* and *P. rioparanaibano* (2.3%, Table 4), it was possible to observe that both cytotypes of *P. rivularis*, inhabitants of the São Francisco River, are closer to *P. paranae* than *P. rioparanaibano* (Figure 2).

Deepening the knowledge of the mitogenome control region, called D-loop, can play a fundamental role in understanding the evolutionary history of the *Astyanax* and *Psalidodon* genera. In this work, we observed that the size variation between different *Astyanax* / *Psalidodon* mitogenomes occurs mainly due to the extension of the D-loop. Neglecting this region in the reconstruction of mitogenomes can result in a valuable loss of information, since in addition to the variation in size, we found a repetitive sequence of 35bp in 9 of the 10 mitogenomes studied (table 3).

*D. giton* was the only species that did not present the repetitive sequence in the D-loop. Plus to occupying a sister group position in the recuperated phylogeny (Figure 2) when describing the mitogenome of the species, Barreto et al [32] observed in their phylogeny of Characidae that the species *Grundulus bogotensis* was closer to the *Astyanax/Psalidodon* group than *D. giton*. Therefore, we not only confirm *D. giton* as a species outside the genus *Astyanax* as suggested by the taxonomic review [4], as the repetitive sequence found in the D-loop may correspond to a synapomorphy absent in the *Deuterodon* group. To confirm this it would be necessary to reassemble the genome of *D. giton* under our methodology and complementary studies need to be done on species of the *Deuterodon* group to clarify this issue.

The methodology used in the reconstruction of the mitochondrial genome proved to be satisfactory and able to access the length of this type of genome, plus the composition and nature of the D-loop, solving possible gaps in previous methodologies [31] [32] [33]. Besides, the study of the complete mitochondrial genome proves to be a tool with the potential to solve taxonomic problems and to help to understand the evolutionary relationships in species complexes, such as *A. bimaculatus*, *P. fasciatus* and *P. scabripinnis*.

## Acknowledgments

Authors thank Henrique Peluzio for his support on the UFV cluster.

The Galaxy server that was used for some calculations is in part funded by Collaborative Research Centre 992 Medical Epigenetics (DFG grant SFB 992/1 2012) and German Federal Ministry of Education and Research (BMBF grants 031 A538A/A538C RBC, 031L0101B/031L0101C de.NBI-epi, 031L0106 de.STAIR (de.NBI)).

